# Sleepless and Desynchronized: Impaired Inter Trial Phase Coherence of Steady-State Potentials Following Sleep Deprivation

**DOI:** 10.1101/471730

**Authors:** M Eidelman-Rothman, E Ben-Simon, D Freche, A Keil, T Hendler, N Levit-Binnun

## Abstract

Sleep loss has detrimental effects on cognitive and emotional functioning. These impairments have been associated with alterations in EEG measures of power spectrum and event-related potentials, however the impact of sleep loss on inter trial phase coherence (ITPC), a measure of phase consistency over experimental trials, remains mostly unknown. ITPC is thought to reflect the ability of the neural response to temporally synchronize with relevant events, thus optimizing information processing.

In the current study we investigated the effects of sleep deprivation on information processing by evaluating the phase consistency of steady-state visual evoked potentials (ssVEPs) as well as amplitude-based measures of ssVEP, obtained from a group of 18 healthy individuals following 24 hours of total sleep deprivation and after a night of habitual sleep. An ssVEP task was utilized, which included the presentation of dots flickering at 7.5 Hz, along with a cognitive-emotional task. Our results show that ITPC is significantly reduced under sleep deprivation relative to habitual sleep. Interestingly, decreased ITPC under sleep deprivation was associated with decreased behavioral performance in the psychomotor vigilance task (PVT), a validate measure of reduced vigilance following lack of sleep.

The results suggest that the capability of the brain to synchronize with rhythmic stimuli is disrupted without sleep. Thus, decreased ITPC may represent an objective and mechanistic measure of sleep loss, allowing future work to study the relation between brain-world synchrony and the specific functional impairments associated with sleep deprivation.

## 2. Introduction

Sleep is a ubiquitous phenomenon, essential for well-being and for optimal behavioral performance. At the neural level, it is hypothesized to have a functional role in various restorative processes, including synaptic plasticity, metabolic upkeep and the balance between excitation and inhibition in neural circuits (Meisel et al., 2013; Tononi and Cirelli, 2006). Accordingly, sleep loss adversely affects cognitive and emotional functioning (Krause et al., 2017). It is associated with decreased performance in cognitive tasks, (Killgore, 2010), overall slowing of responses (Lim and Dinges, 2008), disturbed mood, and impaired emotional processing (Kahn et al., 2013; Pilcher and Huffcutt, 1996; Walker and van Der Helm, 2009).

Several neurophysiological mechanisms have been proposed to underlie the behavioral impairments that accompany sleep loss, including altered functional connectivity patterns (e.g., (Chengyang et al., 2017; De Havas et al., 2012; Lei et al., 2015; Verweij et al., 2014) and reduced event-related potentials/fields features (ERPs/ERFs) associated with the processing of sensory stimuli as well as task-related attention (i.e., N1,P1, P3, e.g., Boonstra et al., 2005; Hoedlmoser et al., 2011; Lee et al., 2003; Morris et al., 1992). For instance, an EEG study (Hoedlmoser et al., 2011) examined the P100 ERP component following sleep deprivation using a visual attention task, known to be sensitive to changes in vigilance (psychomotor vigilance task ;PVT; Drummond et al., 2005). The study demonstrated a progressive decrease in P100 with accrued time awake. The authors further found decreased inter trial phase locking of neural oscillations in the delta and theta frequency ranges during the PVT, which was positively correlated with self-reported sleepiness. This suggests that sleep loss impacts the temporal synchronization of the neural response to external stimuli across several frequency bands.

Neural oscillations reflect periodic fluctuations of excitability in local groups of neurons, as measured noninvasively in humans using EEG\MEG (Buzsáki and Draguhn, 2004; Buzsáki and Watson, 2012; Cohen, 2017; Thut et al., 2012). Fluctuations between high and low cortical excitability states are represented by the phase of neural oscillations, which is the time-varying angle of the oscillatory signal (Buzsáki, 2010; Buzsáki and Watson, 2012; Klimesch et al., 2007; Lakatos et al., 2007, 2008; Thut et al., 2012). In humans, inter trial phase locking of neural oscillations, also referred to as inter trial phase coherence (ITPC), is a measure of phase consistency of the neural response over experimental trials (van Diepen and Mazaheri, 2018).

Increased phase consistency has been associated with enhanced cognitive performance, including enhanced visual perception (Hanslmayr et al., 2005), attention (Ding et al., 2005; Kim et al., 2007) and memory performance (Fell et al., 2008; Klimesch et al., 2004), while decreased consistency has been observed in a number of disorders, such as dyslexia (Hämäläinen et al., 2012), attention-deficit/hyperactivity disorder (ADHD; McLoughlin et al., 2014) and schizophrenia (Teale et al., 2008). Since ITPC reflects the degree of temporal synchronization of the neural response with task-related sensory stimuli, it offers a novel way to examine the neural mechanisms that underlie behavioral impairments following sleep loss, beyond alterations in amplitude or connectivity.

Currently very little is known about the impact of sleep loss on ITPC and specifically, whether the measure of ITPC would demonstrate any sleep sensitivity in paradigms that overtly trigger rhythmic neural activity, such as steady-state visually evoked potentials (ssVEP). ssVEPs are continuous EEG responses, generated by delivering a rhythmic stimulus at a known frequency rate (e.g., flickering dots; Norcia et al., 2015). The ssVEPs are typically recorded in occipital electrodes and are measured as oscillatory waveforms, stable in phase and amplitude (Regan, 1966), with the same frequency peak as the delivered stimulus. Thus, ssVEPs are relatively narrow band oscillations and are advantageous in their excellent signal to noise ratio, which is essential for a reliable phase coherence estimation (Tallon-Baudry et al., 1996; van Diepen and Mazaheri, 2018). In addition, ssVEPs can be detected on a single trial level (Vialatte et al., 2010), which empowers their relevance for behavioral performance.

In the present study we investigated the ITPC of steady-state visual evoked responses (ssVEP), that were recorded under total sleep deprivation and after habitual sleep, in the same participants during the same circadian time. We examined ITPC along with amplitude-based measures and with behavioral performance in the ssVEP task. We further recorded behavioral responses in the psychomotor vigilance task in each sleep condition during the same circadian time, to examine associated changes in vigilance following sleep loss.

**Our main hypothesis is that ITPC during the ssVEP task will be reduced in the sleep deprivation condition compared to habitual sleep, and that this reduction will be accompanied by a decrease in task performance as well as in measures of vigilance.**

## 3. Methods

### 3.1. Participants and Experimental Design

Participants and experimental design were as described in Ben-Simon et al., 2015. Briefly, 18 healthy adults (age range, 23–32 years; mean, 26.8 ± 3 years; 10 females) participated in two experimental sessions each, in a repeated-measures crossover design: after a night of normal sleep (sleep-rested condition, sleep rested) and following 24 h of supervised sleep deprivation (sleep deprivation). EEG was recorded at ∼8:00 A.M. (± 30 min) of the following morning of each session while participants performed a steady-state visual evoked potential (ssVEP) task (see section 3.2). Experimental sessions were separated by a mean of 13.8 d with the order of the sleep-rested and sleep-deprived sessions counterbalanced across participants. Prior to study participation, normal sleep-wake patterns were validated using actigraphy (movement sensor sensitive to wake–sleep states; Fitbit) and subjective sleep logs. During the sleep-rested night, overnight sleep parameters were validated using an ambulatory sleep device (WatchPAT-100; Itamar Medical). Behavioral measures of vigilance were obtained every 2 h during the sleep deprivation night (from 11:00 P.M. to 7:00 A.M.) and in the morning of the sleep-rested session (8:00 A.M. ±30min). These included the Hebrew version of the Stanford Sleepiness Scale questionnaire (SSS; Hoddes et al., 1973) as well as the Psychomotor Vigilance Task (PVT; Drummond et al., 2005), as described in Ben-Simon et al., 2015. Following artifact rejection (as described below), one participant was excluded from further analysis due to insufficient number of artifact-free trials.

### 3.2. ssVEP Task

ssVEPs were elicited using random-dot kinematograms that consisted of randomly moving dots flickering at a rapid rate (7.5 Hz; Ben-Simon et al., 2015). During the presentation of the dots participants were engaged in a visual task aimed at detecting very short intervals of coherently moving dots, while ignoring task-irrelevant neutral or affective distracting pictures, presented at the background of the dots (Deweese et al., 2014; Müller et al., 2007; Figure 1). Prior studies have demonstrated that such competition between the main flickering stimuli and task-irrelevant distractors can be quantified as an attenuation of task-evoked processing, predominantly evident in visual regions (Ben-Simon et al., 2015; Deweese et al., 2014; Müller et al., 2007). It should be mentioned that this aspect of the experimental design is irrelevant for the current research question. Nevertheless, no behavioral effect was found for the presented distractors (Ben-Simon et al., 2015).

**Figure 1.**
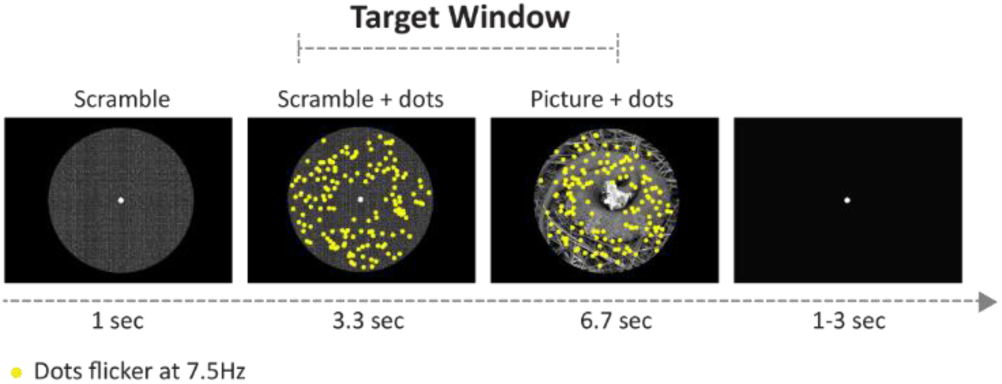
Experimental trial design. Each trial started with 1 s presentation of a scrambled picture, followed by the appearance of flickering dots (at a rate of 7.5 Hz) for 3.3 s. Consequently, a positive, negative, or neutral picture appeared for 6.7 s at the background of the dots. Targets were rare intervals of coherent motion of the dots that could only occur between 1.17 and 7 s after stimulus onset (marked target window). Each trial lasted 10 s, with a variable 1-3 s inter trial interval.

#### 3.2.1. Picture Stimuli

Distractor pictures were divided into three valence categories, positive, negative and neutral (for details see Ben-Simon et al 2015). Each category included 30 pictures, totaling 90 pictures, selected from the IAPS (Lang et al., 1997), with additional images selected from the public domain to complete balanced human and animal picture categories. All stimuli were grayscale pictures and were controlled for visual complexity (measured as.jpeg size) and matched for luminance using scripts from the MATLAB image processing toolbox (picture stimuli were circular in nature and were cropped and adjusted such that the defining element of each picture was positioned at the center of a circle (see Figure 1).

### 3.3. Experimental Trial

Each trial began with a 1 s presentation of a stimulus image with individual pixels scrambled to avoid contamination of the ssVEP with transient responses to the luminance gradient created by stimulus onset (Ben-Simon et al., 2015). Next, a total of 150 yellow dots (each 0.3° x 0.3° of visual angle) were superimposed on the scrambled image for 3.3 s. The scrambled picture was then replaced by a positive, neutral, or negative image that remained on the screen for the remaining duration of the trial (6.7 s). The flickering dots were distributed randomly across pictures and remained inside the circle (6.9° visual angle) at all times. The yellow dots were “on” for four frames and “off” for four frames. All dots remained in continuous motion throughout the trial, and each dot changed its position by 0.04° in a random direction with every ssVEP cycle (i.e., 7.5 times/s).

In a random subset of 50% of the trials, all dots moved coherently in the same direction (target), and participants were instructed to respond to coherent motion events with a mouse click, as quickly and as accurately as possible. Coherent motion of the targets occurred in one of four diagonal directions (45°, 135°, 225°, and 315°) at random. In an effort to produce a difficult and demanding perceptual detection task, coherent motion lasted for only four successive cycles of 7.5 Hz (i.e., 533.33 ms). Each trial lasted 10,000 ms, with inter stimulus intervals varying randomly between 1,000 and 3,000 ms, during which a white fixation dot was presented at the center of the screen (see Figure 1). Stimuli were presented centrally on an LED monitor, set at a resolution of 1024 × 768 with a refresh rate of 60 frames/s (i.e., 16.66 ms refresh interval).

### 3.4. EEG Data Recording and Preprocessing

The EEG signal was recorded from the scalp using the BrainAmp-MR EEG amplifier (Brain Products) and the BrainCap electrode cap with sintered Ag/AgCl ring electrodes providing 30 EEG channels, one EKG channel, and one EOG channel (Falk Minow Services). The reference electrode was between Fz and Cz. Raw EEG was sampled at 500 Hz and recorded using the Brain Vision Recorder software (Brain Products) with scalp impedance for each electrode kept below 20 kΩ. Signal preprocessing was carried out using MATLAB (The MathWorks) and functions from the EEGlab toolbox (Delorme and Makeig, 2004). The continuous data were bandpass filtered offline in the 1-40 Hz range (Hamming windowed sinc FIR filter). Subsequently, the data was segmented into 16 s epochs (−4 s before and 12 s after dots onset) and segments with amplitudes exceeding 100 µV were excluded from further analysis. Blinks and eye movement artifacts were subsequently removed from the data utilizing the Second Order Blind Identification (SOBI) (Belouchrani et al., 1997) Independent Component Analysis (ICA) algorithm, implemented in FieldTrip (Oostenveld et al., 2010). Following these preprocessing steps, the overall average trial retention rate was 80.23 trials, with trial counts not significantly different between conditions (positive, 26.73 trials; neutral, 27.09 trials; negative, 26.47 trials on average; P > 0.1, paired t-tests) or experimental sessions (sleep-rested, 84.09; sleep-deprived, 84.66 trials on average, P > 0.1, paired t-test).

### 3.5. ssVEP Analysis

The analysis was focused on occipital electrodes (O1, Oz, O2), where the greatest overall ssVEP amplitudes have been observed (Ben-Simon et al., 2015). The effect of emotional distractor on ssVEP was analyzed and reported in previous work with the same experimental group (Ben-Simon et al., 2015) and the current analysis was performed on the entire trial pool, including all distractor types.

#### 3.5.1. ssVEP Signal Evaluation

To visualize and evaluate the ssVEP signal, the pre-processed time-domain EEG data was averaged across trials, electrodes, sleep conditions and subjects. In addition, the frequency content of the data was estimated by subjecting each trial to a fast Fourier transform (FFT) analysis (Figure 2) to verify that the flickering dots reliably evoked steady-state responses in the expected stimulation frequency.

**Figure 2.**
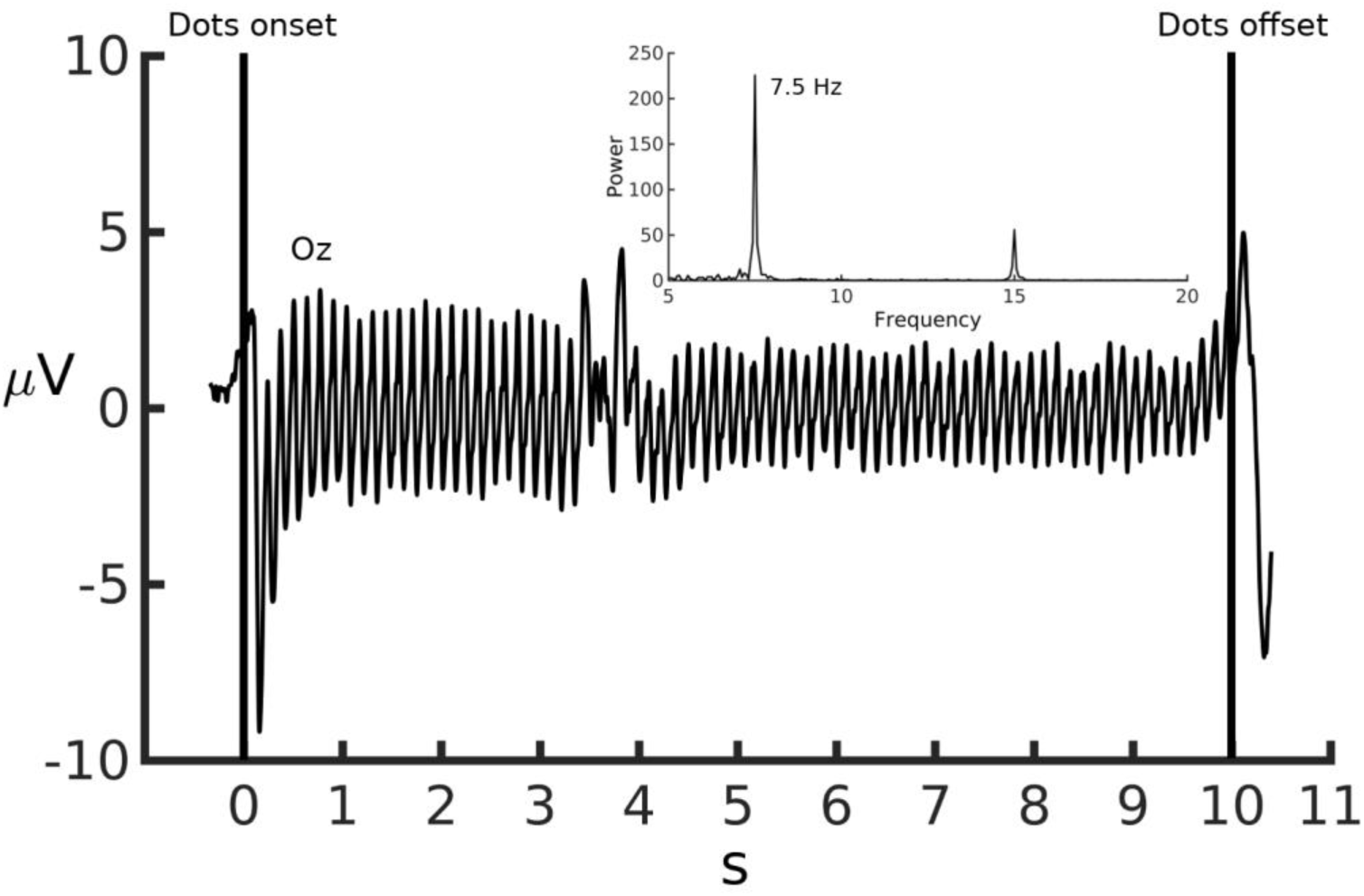
Illustration of the steady-state visual evoked signal. Time domain data from Oz electrode and FFT power spectrum (inset) averaged across trials, sleep conditions and across subjects. A reliable peak in the power spectrum is evident at the stimulation frequency (7.5 Hz) and its second harmonic (15 Hz).

#### 3.5.2. ssVEP Inter Trial Phase Coherence Analysis

The pre-processed segmented EEG data was filtered around the stimulation frequency (7.5 Hz ± 0.5) using Hamming windowed sinc FIR filter (order 826, transition bandwidth 2 Hz, cutoff frequencies 6-9 Hz, stopband attenuation -53 dB). For inter trial phase coherence (ITPC) analysis, the filtered data was subjected to Hilbert transform. Phase-locking values (PLVs; i.e., the resultant vector length) were calculated for each time point using the following formula, as previously described (Lachaux et al., 1999; Sharon and Nir, 2017; van Diepen and Mazaheri, 2018):

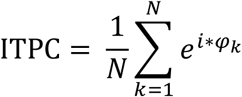

where N is the number of trials and *φ*_k_ is the angle of the signal relative to the stimulus, in radians. PLVs were calculated as the absolute value of the ITPC, yielding values between 0 (high phase variability across trials) and 1 (uniformity of phase across trials). The PLVs were averaged across the three occipital electrodes (O1, O2, Oz) for each participant, and group statistics comparing between sleep conditions was carried out using paired t-tests (implemented in MATLAB). Paired t-tests were conducted in 1000 ms time windows along the experimental trial, with False Discovery Rate (FDR) correction for multiple comparisons (Benjamini and Hochberg, 1995).

#### 3.5.3. ssVEP Amplitude-Based Measures Analysis

Three measures of ssVEP amplitude were calculated: evoked amplitude, total amplitude and baseline-corrected total amplitude. For evoked amplitude analysis, the Hilbert transform was applied to the averaged time-domain filtered-data (see section 3.5.2), while the total amplitude was obtained by applying the Hilbert transform to each trial, thus retaining activity that is not phase-locked to the stimulus. The evoked and total amplitude at each time point was extracted as the absolute value of the Hilbert transformed analytic signal. The baseline-corrected total activity was calculated by subtracting the mean amplitude of a 0.5 s pre-stimulus period, between 1.5 s to 2 s prior to dots onset, from each time point along the trial. This measure was used to evaluate differences between conditions during the steady-state response time period, that are not related to differences in baseline activity. Group statistics comparing the three amplitude-based measures between sleep conditions was conducted similarly to the PLV analysis as described in section 3.5.2.

#### 3.5.4. Ongoing Activity Analysis

Decreased phase synchronization during processing of external events is hypothesized to be associated with elevated background noise (Krystal et al., 2017) such as spontaneous activity. We therefore examined ongoing brain activity over a broad frequency range, in sleep rested vs. sleep deprivation. To this end, spectral analysis was performed on each trial in each occipital electrode. Spectral power was estimated in the 1-25 Hz frequency range using the short-time Fourier transform with a sliding hamming window of 1024 samples length and 1023 overlapped samples. The obtained spectrograms were averaged across trials, occipital electrodes and across subjects. Subsequently, the averaged spectral power within the delta (1-4 Hz), theta (4-8 Hz) and alpha (8-12 Hz) frequency ranges, known to be affected by sleep deprivation (Bernardi et al., 2015; Hoedlmoser et al., 2011; Nir et al., 2017), was compared between sleep conditions. For each frequency band, two different time windows were tested: a baseline window (between -1 s and -3 s before dots onset) and the steady-state response window. Two values were consequently calculated for each frequency (delta, theta, alpha) in each sleep condition (sleep rested/sleep deprivation), per subject, by averaging the spectral power over all time points within each time window. Spectral power differences between conditions were tested using paired t-test.

### 3.6. Correlation Between Amplitude-Based Measures and ITPC

ITPC differences between experimental conditions have been shown to be affected by corresponding differences in amplitude (van Diepen and Mazaheri, 2018). We therefore tested the relation between the differences in the ssVEP amplitude-based measures (evoked amplitude, total amplitude, baseline-corrected total amplitude) and in ITPC as a result of sleep deprivation. A single value was first calculated for each measure (amplitude/phase) in each sleep condition (sleep rested/sleep deprivation) per subject, by averaging the amplitudes/PLVs over all time points along the steady state visual evoked response (i.e., from dots onset to dots offset). The difference between conditions (sleep rested minus sleep deprivation) in mean PLV was then correlated with the corresponding difference in ssVEP amplitude, using Spearman correlation.

### 3.7. Correlation Between ITPC and Behavioral Measures

As reported in Ben-simon et al., 2015, sleep deprivation resulted in increased subjective and objective assessments of sleepiness (the Stanford Sleepiness Scale questionnaire and the psychomotor vigilance task, respectively) and in decreased performance in the ssVEP task. The association between sleep deprivation-related alterations in ITPC and in behavioral measures was tested by correlating the mean PLV (as calculated in section 3.6) with each behavioral measure under sleep deprivation, as well as the difference in PLV between conditions (sleep rested minus sleep deprivation) with the corresponding difference in the behavioral measures, using Spearman correlation.

### 3.8. Correlation Between Ongoing Activity and ITPC

To test whether increased ongoing activity is related to decreased ITPC under sleep deprivation, the mean PLV (as calculated in section 3.6) was correlated with the mean ongoing activity during the steady-state evoked response period (as calculated in section 3.5.4) in each frequency band (delta, theta, alpha) under sleep deprivation, using Spearman correlation.

## 4. Results

### 4.1. The ssVEP signal

The steady-state visual evoked signal is illustrated in Figure 2. A reliable peak at the stimulation frequency (7.5 Hz) and its second harmonic (15 Hz) is clearly seen in the FFT power spectrum (Figure 2 inset).

### 4.2. Inter trial Phase Coherence of ssVEPs is Decreased in Sleep Deprivation

Our inter trial phase-locking analysis revealed a significant reduction in PLVs in the sleep deprived compared to the sleep rested condition throughout the steady state evoked response period (i.e., from dots onset to dots offset, excluding the time window between 8 to 9 s following dots onset; Figure 3a). t values ranged from t(16) = 3.78 to t(16) = 5.65, all p values ≤ 0.003, FDR corrected, and effect ranged sizes from 0.74 to 1.37 (Cohen’s d), across all time windows along the ssVEP period. This was the case for the vast majority of the participants as demonstrated in Figure 3b and 3c. These findings indicate that sleep deprivation is associated with increased variability in the timing of the ssVEP responses, which can be seen on the individual subject level, suggesting that decreased ITPC may be used as a reliable indicator of sleep deprivation.

**Figure 3.**
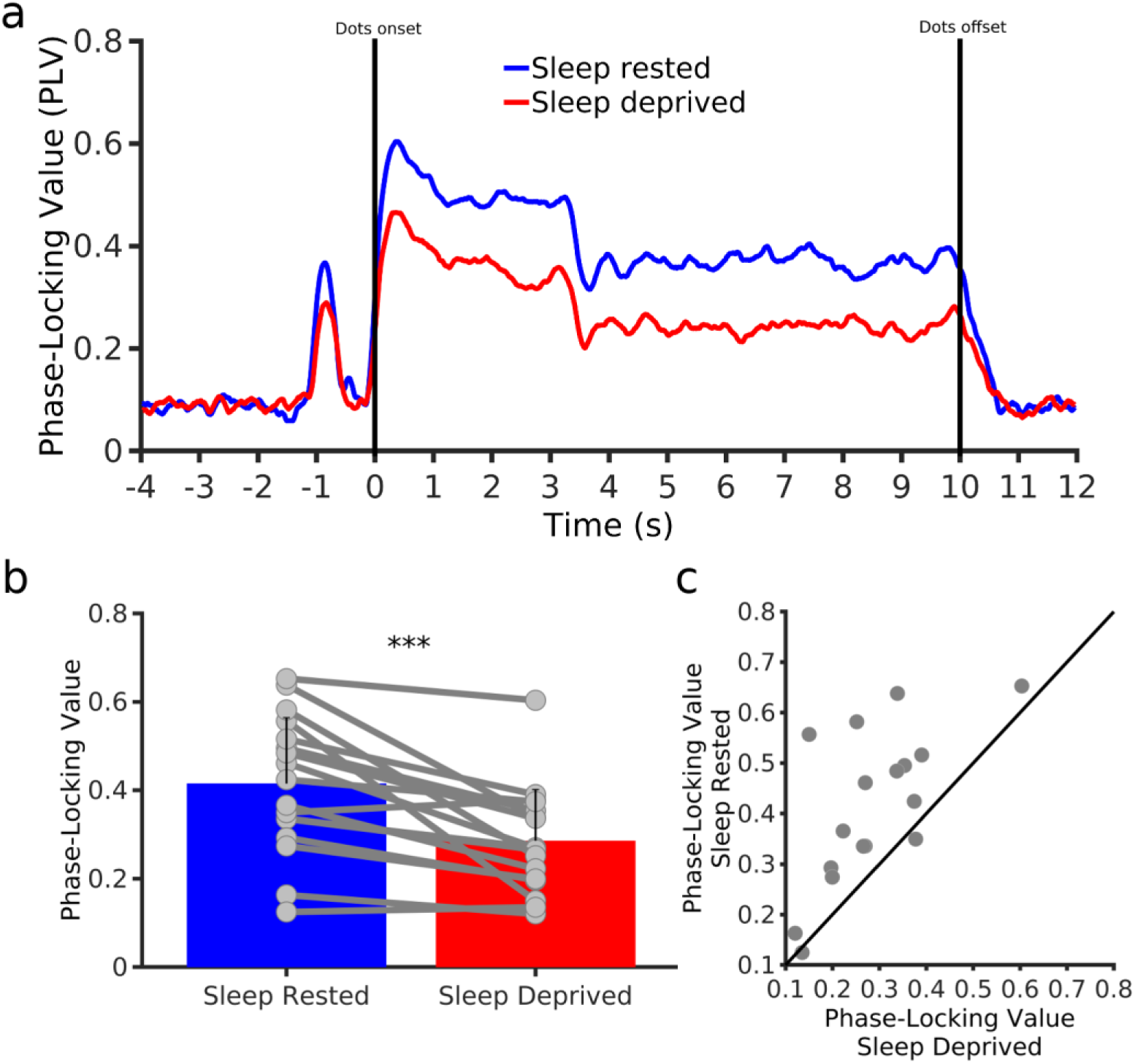
The effect of sleep deprivation on inter trial phase coherence of the steady-state visual evoked response. (a) PLVs calculated around the stimulation frequency (7.5 Hz), averaged across occipital electrodes (O1, O2, Oz), and across subjects. PLVs during the steady-state evoked response period (i.e., from dot onset at time zero to dot offset at 10 s) were significantly lower in the sleep deprived (sleep deprivation) compared to the sleep rested (sleep rested) condition (all p values ≤ 0.003, FDR corrected). (b) Group level (bar graph) and individual level (circles) differences between sleep rested and sleep deprivation in mean PLVs, calculated per individual as the averaged values over all time points within the steady-state evoked response. Error bars represent standard deviations. t(16) = 4.47, p = 0.0004, Cohen’s d = 1.08. (c) Mean PLVs in sleep rested and sleep deprivation plotted against each other. Each circle represents one subject. As indicated by the position of most circles above the diagonal line (i.e., equality of PLVs in sleep deprivation and sleep rested), decreased inter trial phase coherence is evident at the individual level for almost the entire sample.

#### 4.2.1. ITPC is a More Sensitive Measure for Sleep Deprivation than Amplitude-Based Measures

To evaluate the effect of sleep deprivation on ssVEP amplitude, differences between sleep conditions were examined using three amplitude-based measures: evoked amplitude, total amplitude and the baseline-corrected total amplitude. Group averaged activity along the experimental trial is presented for each measure in Figure 4a.

**Figure 4.**
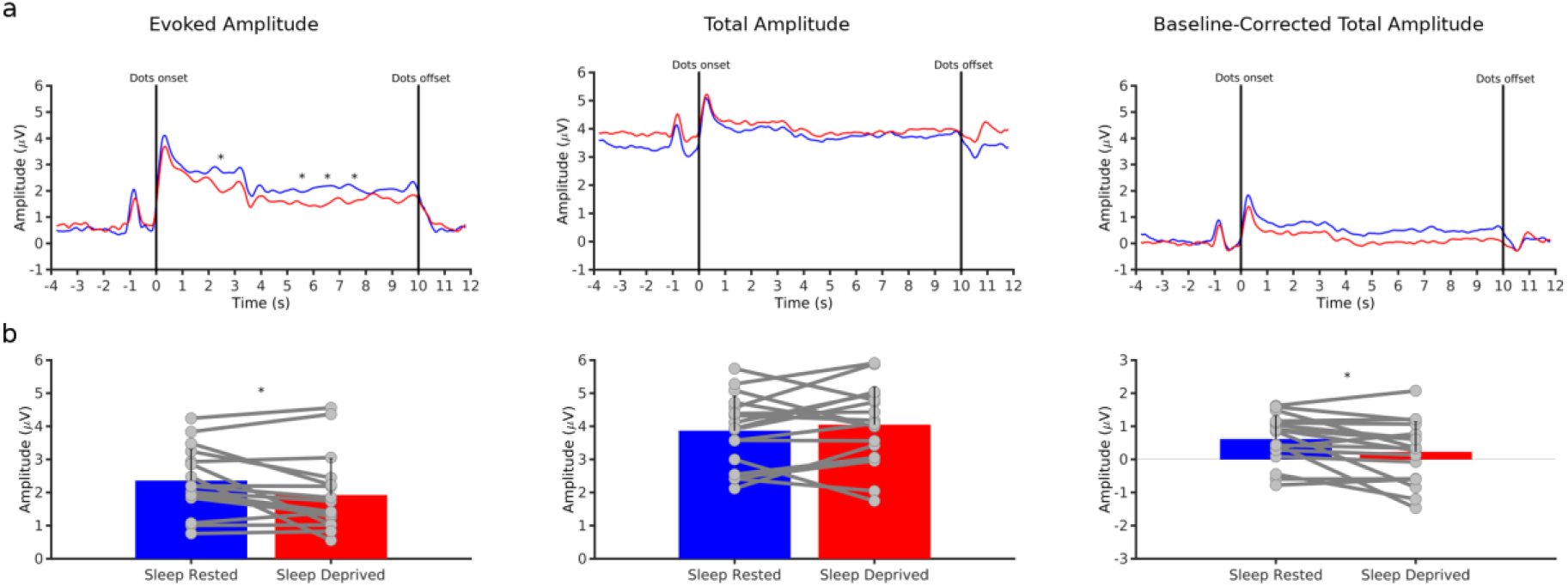
The effect of sleep deprivation on amplitude-based measures of the steady-state visual evoked response. (a) Amplitude measures calculated around the stimulation frequency (7.5 Hz), averaged across occipital electrodes (O1, O2, Oz), and across subjects. Evoked amplitude (left panel) during the steady-state response period (i.e., from dot onset to dot offset) was significantly lower in the sleep deprived compared to the sleep rested condition in specific time windows (marked with asterisks), starting from 2s following dots onset (all p values ≤ 0.048, FDR corrected). Differences between conditions in total amplitude (with and without baseline correction) were not significant in all tested time windows (middle and right panels). (b) Group level (bar graph) and individual level (circles) differences between sleep rested and sleep deprivation in mean amplitude (evoked, total and baseline-corrected total), calculated per individual as the averaged values over all time points within the steady-state evoked response. Error bars represent standard deviations. Significant differences were found for evoked amplitude (t(16) = 2.5, p = 0.023, Cohen’s d = 0.6) and for the baseline-corrected total amplitude (t(16) = 2.38, p = 0.03, Cohen’s d = 0.57) but not for total amplitude with no baseline correction (t(16) = -0.9, p = 0.37, Cohen’s d = -0.21),

ssVEP evoked amplitude was higher in the sleep rested compared to the sleep deprivation condition (left panel). These differences were statistically significant in specific time windows along the steady-state evoked response period. T values ranged from t(16) = 2.82 to t(16) = 3.49; all p values ≤ 0.048, FDR corrected, and effect sizes ranged from 0.22 to 0.84 (Cohen’s d).

The total ssVEP amplitude in sleep deprivation was higher compared to the sleep rested condition throughout the experimental trial (middle), however this was not statistically significant (t(16) = -0.41 to t(16) = -1.49; all p values ≥ 0.35, FDR, Cohen’s d = -0.1 to -0.36 for time windows within the ssVEP period). Following baseline correction, sleep rested total amplitude was higher compared to sleep deprivation during the ssVEP period (right panel), however not significantly (t(16) = 1.6 to t(16) = 2.81; all p values ≥ 0.085, FDR corrected, Cohen’s d = 0.4 to 0.68).

For each amplitude measure, the time-averaged activity within the ssVEP period (from dots onset to dots offset; as calculated in section 3.6) was compared between sleep conditions (Figure 4b), similarly to the PLV measure presented in Figure 3b. Inspection of the PLV vs. the amplitude-based measures in this analysis clearly demonstrates that ITPC discriminates more reliably between the sleep rested and sleep deprivation conditions, both at the individual subject level and at the group level. This is quantitatively demonstrated by the larger effect size measured for changes in PLV compared to the amplitude-based measures (Cohen’s d values = 1.08, 0.6, -0.21 and 0.57, for PLV, evoked amplitude, total amplitude and baseline-corrected total amplitude, respectively). In light of its higher sensitivity to sleep deprivation, ITPC was used in the correlation analyses described in the following section, rather than amplitude-based measures.

#### 4.2.2. Correlation Between ITPC and ssVEP Amplitude-Based Measures

It has been shown that ITPC differences between experimental conditions may be affected by corresponding differences in the amplitude of the signal (van Diepen and Mazaheri, 2018). Therefore, in addition to the evaluation of differences in amplitude-based measures between the sleep rested and sleep deprived conditions as described in section 4.2.1, we correlated the differences in mean PLVs and mean amplitudes between sleep conditions. A significant positive correlation was found for differences in PLVs and in evoked amplitudes (r = 0.68, p = 0.0031) and baseline-corrected total amplitudes (r = 0.7, p = 0.021; supplementary Figure S1), indicating that a decrease in ITPC was associated with a corresponding decrease in these amplitude measures. The correlation between differences in PLVs and in the non baseline-corrected total amplitudes were not significant (r = 0.22, p = 0.38; supplementary Figure S1).

#### 4.2.3. Ongoing Activity is Increased in Sleep Deprivation

Group averaged time-frequency representations of the sleep rested and sleep deprived conditions data are presented in Figure 5. The steady state response in 7.5 Hz and its second harmonic at 15 Hz are seen in the two conditions, as well as the expected attenuation (relative to baseline) in alpha range (8-12) activity throughout the steady-state response period (Keitel et al., 2019). As demonstrated in the difference plot of the two spectra (sleep rested minus sleep deprived; lower panel), the spectrum in the sleep deprived condition is characterized by increased low-frequency power, in the alpha, theta and delta frequency ranges compared to the sleep rested condition. These differences were statistically significant, both in the baseline time window (t(16)_alpha_ = -2.74, p(16)_alpha_ = 0.014; t(16)_theta_ = -2.86, p_theta_ = 0.011; t(16)_delta_ = -2.64, p_delta_ = 0.017) and in the steady-state response time window ((16)_alpha_ = -3.36, p_alpha_ = 0.004; t(16)_theta_ = -3.65, p_theta_ = 0.002; t(16)_delta_ = -3.77, p_delta_ = 0.0016; see supplementary Figure S2).

**Figure 5.**
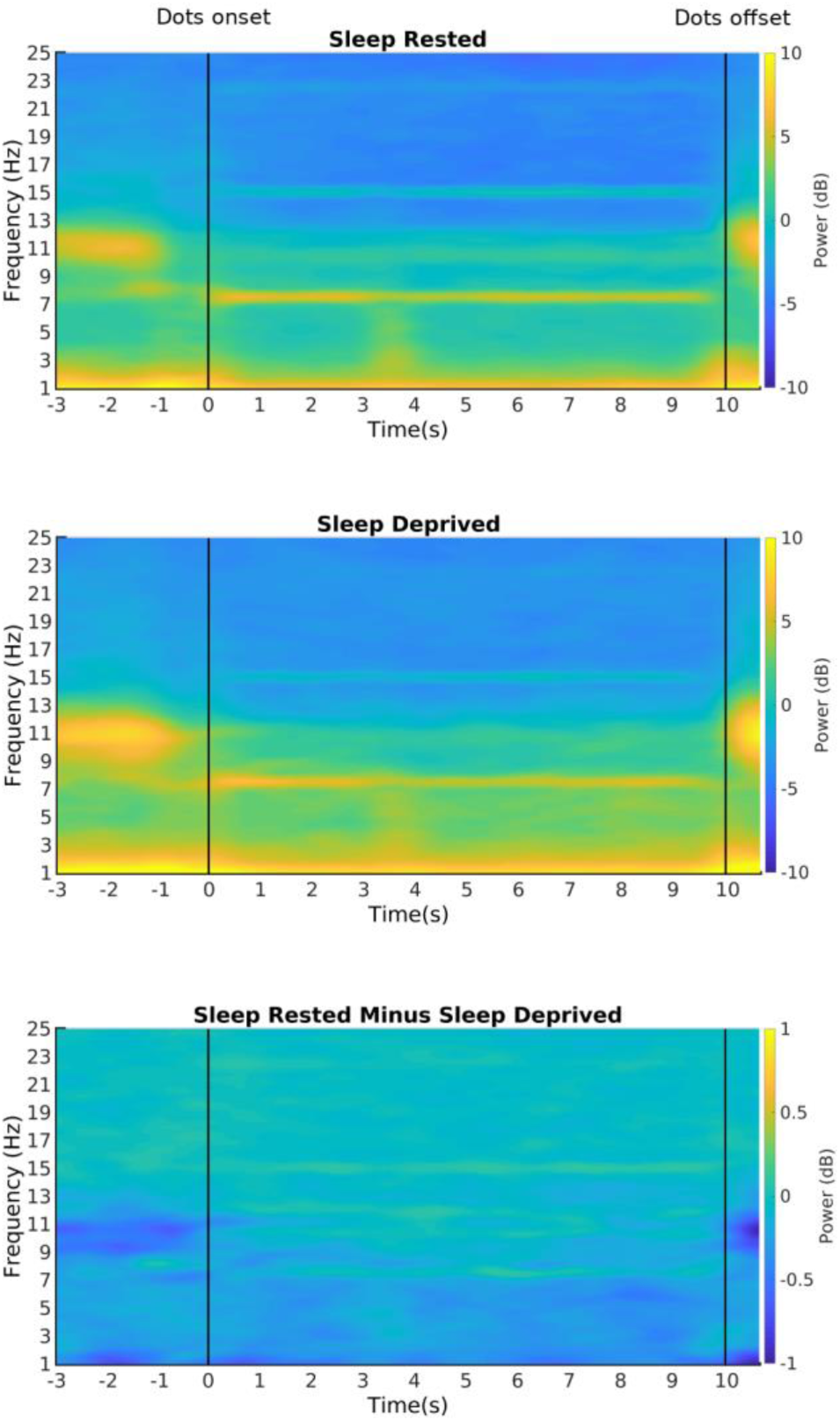
Time-frequency representation of ongoing activity. Spectrograms were averaged across occipital electrodes (O1, O2, Oz) and across subjects. The steady state response in 7.5 z and its second harmonic at 15 Hz is seen in the two sleep conditions. As revealed by the difference plot (lower panel, sleep rested minus sleep deprived), spectral power in the delta, theta and alpha frequency ranges was overall greater in sleep deprivation relative to the sleep condition.

### 4.3. Relation Between ITPC and Behavioral Measures following sleep loss

To further evaluate the measure of ITPC in sleep deprivation, we examined the association between phase-locking values (mean PLV as calculated in section 3.6) and self-reported sleepiness as well as task performance, under sleep deprivation. In addition, we tested the relations between the observed reduction in phase coherence and the changes in the different behavioral measures, from sleep rested to sleep deprivation. The latter was tested by correlating the difference between the mean PLV in sleep rested and sleep deprivation with the corresponding differences in the scores of the sleepiness scale (level of fatigue), the PVT (number of lapses) and the ssVEP task (accuracy).

A significant correlation was found between decreased PLVs in the sleep deprived condition and increased number of lapses in the PVT task (r = -0.5, p = 0.038), suggesting that lower phase locking values are associated with impaired performance in the PVT task. In order to test the specificity of this effect to sleep deprivation, the correlation analysis was repeated for the sleep rested condition, however, no correlation effect was found (r = 0.12, p = 0.63). This suggests that the association between these measures is specific to and driven by sleep loss. In addition, the correlation between decreased ITPC and impaired task performance remained significant also when statistically controlling for evoked amplitude levels (partial correlation analysis; r = -0.5, p = 0.028), which as was shown above, found to be significantly affected by sleep deprivation in several time points along the experimental timeline (Figure 4a left panel). This finding points at the possibility that similar to the PVT, known for its high sensitivity to sleep deprivation (Drummond et al., 2005), PLV may serve as a reliable measure for assessing the functional impairments triggered by sleep loss. All other mentioned correlation analyses were not statistically significant (see supplementary results).

### 4.4. Relation Between ITPC and ongoing activity in sleep deprivation

Decreased phase synchronization during stimuli processing is hypothesized to be related to elevated background noise (Krystal et al., 2017). The relation between ongoing activity and ITPC in sleep deprivation was therefore tested by correlating the mean PLVs (as calculated in section 3.6) with the ongoing activity measured during the steady-state response period (as calculated in section 3.5.4) in the delta, theta and alpha frequency bands. A significant negative correlation was found between PLVs and theta activity under sleep deprivation (r = -0.55, p = 0.024). In order to test the specificity of this effect to sleep deprivation, the correlation analysis was repeated for the sleep rested condition, however, no correlation effect was found (r = -0.35, p = 0.16). This suggests that the relation between these measures is specific to and driven by sleep loss. As was the case for the association between PLV and the psychomotor vigilance task (section 4.3), the correlation between decreased ITPC and theta activity during the steady-state response period remained significant also when statistically controlling for evoked amplitude levels (partial correlation analysis; r = -0.76, p = 0.0006). The correlations between PLVs and ongoing alpha and delta activity were not significant (r = -0.46, p = 0.06 and r = -0.4, p = 0.11 for alpha and delta respectively).

## 5. Discussion

Our findings demonstrate that inter-trial phase coherence of steady state visual evoked potentials is reduced after one night of total sleep deprivation, in comparison to habitual sleep. Alterations in ssVEP amplitude-based measures were also found under sleep deprivation, however ITPC showed superior sensitivity to sleep loss relative to the amplitude-based measures. Together, these findings suggest that sleep deprivation disrupts the capability of the neural response to temporally synchronize with external stimuli. Despite identical sensory demands put forward by the ssVEP task, the ability to synchronize to the task in a consistent manner was dependent upon a rested night of sleep. Our findings further demonstrate that sleep deprivation can be reliably assessed by means of ssVEPs paradigms, shown to be sensitive to a wide range of tasks used in cognitive and clinical neuroscience (Norcia et al., 2015; Vialatte et al., 2010; Wieser et al., 2016). These results are in line with prior findings that demonstrate decreased phase consistency in sleep deprivation (Hoedlmoser et al., 2011), as well as studies that demonstrate the impact of fatigue on the amplitude of ssVEPs (Cao et al., 2014; Hoedlmoser et al., 2011).

While both the phase-locking and amplitude measures were affected by sleep deprivation, and although these effects were highly correlated, a comparison between ITPC and amplitude-based measures in our study clearly demonstrated higher sensitivity of ITPC to sleep loss. This was indicated by the larger effect size observed for the changes in ITPC from sleep rested to the sleep deprived condition, which were evident at the individual level across nearly the entire experimental group. The high reliability of ITPC in assessing sleep deprivation was further supported by the significant correlation found between decreased ITPC under sleep deprivation and impaired performance in the psychomotor vigilance tasks, as well as with increased theta power, both are known for their high sensitivity to sleep loss (Drummond et al., 2005; Finelli et al., 2000; Nir et al., 2017; Vyazovskiy and Tobler, 2005). For example, a recent study conducted with neurosurgical patients found elevated theta power under total sleep deprivation, as well as higher theta before the occurrence of cognitive lapses during the performance of a categorization psychomotor vigilance task (Nir et al., 2017). Our findings therefore suggest that ITPC may serve as a reliable, noninvasive indicator of sleep deprivation and associated behavioral impairment.

A desynchronized neural response to external stimuli could reflect impairments in information processing following sleep loss. Interestingly, a recent study demonstrated attenuated ssVEP responses and decreased ITPC during nighttime sleep when stimulating at the alpha (8 and 10 Hz) frequency range (Sharon and Nir, 2017). These findings raise the possibility that the reduced phase locking of ssVEPs around 7.5 Hz observed in our study indexes increased sleep propensity following a sleepless night and generally points to the impact of both sleep and sleep loss on neural responses to external stimuli.

Decreased temporal synchronization with external stimuli has further been suggested to reflect increased background noise, such as spontaneous neural activity (Kashiwase et al., 2012; Krystal et al., 2017). In accordance, previous studies found higher spontaneous activity under sleep deprivation, which also indicated elevated sleep pressure (Bernardi et al., 2015; Nir et al., 2017) and impaired behavioral performance (Bernardi et al., 2015; Hung et al., 2013; Nir et al., 2017). A failure to suppress such background noise was theorized to produce impaired behavioral performance, for example due to reduced precision of information representation(Krystal et al., 2017). Our finding of increased ongoing activity (found in the delta, theta and alpha frequency ranges) following sleep deprivation, may thus represent a state of impaired noise suppression, which may have affected the ability of the neural response to synchronize with presented stimuli. Support for this proposition comes from numerous resting state fMRI studies, suggesting that sleep deprivation triggers a breakdown in network integrity that may negatively affect, and possibly predict, task related responses in this condition (Krause et al., 2017). For example, a graph theoretical analysis applied on a resting state fMRI data obtained from the currently tested cohort showed that under conditions of sleep deprivation, network architecture shifts towards a more random-like organization, leading to impaired functional segregation in regions of the limbic, salience and default mode networks (Ben Simon et al., 2017). These alterations were further associated with impaired task performance elicited by sleep deprivation.

The ability to track dynamic visual stimuli has been demonstrated to have an important role in perception and behavioral performance (Keitel et al., 2016, 2019; Kim et al., 2007). We therefore tested whether the decreased ITPC found in sleep deprivation is associated with impaired behavioral task performance. This was the case as decreased PLVs were associated with impaired performance in the PVT task, suggesting that the inability to synchronize can impair task related behavior and specifically one that requires prolonged attention as the PVT (Doran et al., 2001; Jewett et al., 1999), known as an objective marker of reduced vigilance. This could suggest that PLV is more sensitive to global states of arousal.

Our analysis revealed that both the task-related amplitude and the phase coherence of the ssVEP response were decreased under sleep deprivation, and that this decrease was strongly related among the two measures. This finding raises the possibility that the observed reduction in ITPC results from the weaker task-evoked signal measured in sleep deprivation compared to sleep rested (van Diepen and Mazaheri, 2018). Although the possibility that the ITPC differences arise from amplitude differences cannot be completely ruled out, our findings point towards decreased ITPC as a reliable indicator of sleep deprivation, as demonstrated by its correlation with well established markers for sleep deprivation (PVT and theta power), even when statistically controlling for evoked amplitude levels. In addition, no overall effect of reduced power was found in sleep deprivation across multiple frequency bands as indicated by the increased baseline power of lower frequency bands found in this condition. This suggests that the decrease in power during the steady-state response is task-specific and is most likely related to an impaired ability to temporally synchronize with the dynamic visual stimuli, rather than to a global decrease in power. The enhanced low-frequency power found here along with the decrease in ITPC may reflect a transition of the brain into a sleep state and its prioritization over task-related demands. This suggestion is in line with previous research showing the occurrence of sleep-like activity in the awake, sleep deprived human brain, and its association with behavioral impairments (Bernardi et al., 2015; Hung et al., 2013; Nir et al., 2017).

In summary, our main findings indicate that sleep deprivation decreases ITPC of steady state visual evoked potentials and demonstrate that ITPC is a highly sensitive measure for sleep deprivation. The decrease in inter trial phase-locking suggests that in sleep deprivation, responses from a large population of neurons (combined in the ssVEPs) become less synchronized with external stimuli, affecting the capability of the brain response to follow the stimuli rhythm in a temporally precise manner, which ultimately leads to reduced information processing in sleep deprivation. ITPC may serve as a useful and non-invasive method to investigate the effects of sleep deprivation on brain and behavior in healthy individuals and in conditions associated with disrupted sleep.

## 6. Acknowledgements

This study was supported by the Sagol family foundation and by the Joy Academic Grant Program

