## Supplemental data for "Sleepless and Desynchronized: Impaired Inter Trial Phase Coherence of Steady-State Potentials Following Sleep Deprivation"

### Supplementary material

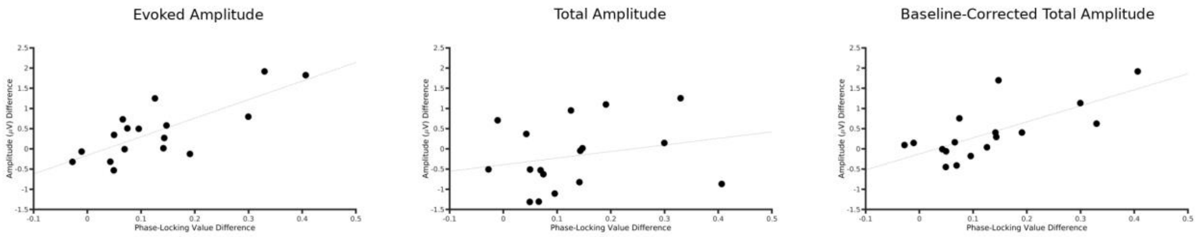


**Figure S1. Correlation between ssVEP amplitude and ITPC.** Correlation between the differences in mean PLVs and in mean amplitudes between sleep conditions (sleep rested minus sleep deprived). Significant correlation was found with evoked (r = 0.68, p = 0.0031; left panel) and for the baseline-corrected total amplitudes (r = 0.7, p = 0.021; right panel), but not with the baseline-corrected total amplitudes (r = 0.22, p = 0.38; middle panel).

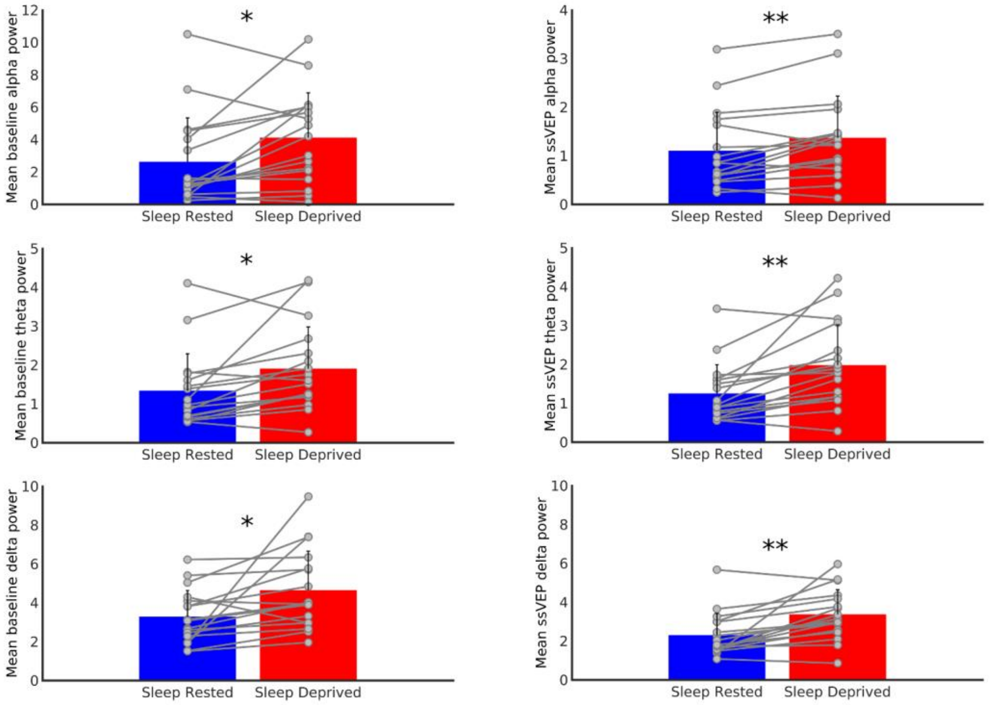


**Figure S2. Ongoing activity in baseline and during the steady-state response period.** Group level (bar graph) and individual level (circles) differences between sleep rested and sleep deprivation in mean ongoing activity in the alpha (8-12 HZ), theta (4-8 Hz) and delta (1-4 Hz) frequency ranges, in the baseline time window ($t(16)_{alpha}$ = -2.74, $p(16)_{alpha}$ = 0.014; $t(16)_{theta}$ = -2.86, $p_{theta}$ = 0.011; $t(16)_{delta}$ = -2.64, $p_{delta}$ = 0.017; left panel) and in the steady-state response time window ($(16)_{alpha}$ = -3.36 , $p_{alpha}$ = 0.004; $t(16)_{theta}$ = -3.65, $p_{theta}$ = 0.002; $t(16)_{delta}$ = -3.77 , $p_{delta}$ = 0.0016; right panel). Error bars represent standard deviations.

### Correlation between ITPC and behavioral measures

Correlations between phase-locking values (mean PLV as calculated in section 3.6) and the scores of the sleepiness scale (level of fatigue) and the ssVEP task (accuracy) under sleep deprivation were not significant (r = 0.0761, p = 0.77; r = 0.051, p = 0.84, respectively). This was also the case for the relations between the observed reduction in phase coherence and the changes in behavioral measures, from sleep rested to sleep deprivation, tested by correlating the difference between the mean PLV in sleep rested and sleep deprivation with the corresponding differences in the scores of the sleepiness scale and the ssVEP task ($r = -0.13$, $p = 0.6$, $r = 0.33$, $p = 0.18$ for , sleepiness scale and the ssVEP task respectively).
